# Canonical and Replicable Multi-Scale Intrinsic Connectivity Networks in 100k+ Resting-State fMRI Datasets

**DOI:** 10.1101/2022.09.03.506487

**Authors:** A. Iraji, Z. Fu, A. Faghiri, M. Duda, J. Chen, S. Rachakonda, T. DeRamus, P. Kochunov, B. M. Adhikari, A. Belger, J.M. Ford, D.H. Mathalon, G.D. Pearlson, S.G. Potkin, A. Preda, J.A. Turner, T.G.M. van Erp, J. R. Bustillo, K. Yang, K. Ishizuka, A. Sawa, K. Hutchison, E. A. Osuch, Jean Theberge, C. Abbott, B.A. Mueller, D. Zhi, C. Zhuo, S. Liu, Y. Xu, M. Salman, J. Liu, Y. Du, J. Sui, T. Adali, V.D. Calhoun

**Author notes:** Correspondence address to (Armin Iraji), (Vince Calhoun).

## Abstract

Resting-state functional magnetic resonance imaging (rsfMRI) has shown considerable promise for improving our understanding of brain function and characterizing various mental and cognitive states in the healthy and disordered brain. However, the lack of accurate and precise estimations of comparable functional patterns across datasets, individuals, and ever-changing brain states in a way that captures both individual variation and inter-subject correspondence limits the clinical utility of rsfMRI and its application to single-subject analyses.

We posit that using reliable network templates and advanced group-informed network estimation approaches to accurately and precisely obtain individualized (dynamic) networks that retain cross-subject correspondence while maintaining subject-specific information is one potential solution to overcome the aforementioned barrier when considering cross-study comparability, independence of subject-level estimates, the limited data available in single studies, and the low signal-to-noise ratio (SNR) of rsfMRI.

Toward this goal, we first obtained a reliable and replicable network template. We combined rsfMRI data of over 100k individuals across private and public datasets and selected around 58k that meet quality control (QC) criteria. We then applied multi-model-order independent component analysis (ICA) and subsampling to obtain reliable canonical intrinsic connectivity networks (ICNs) across multiple spatial scales. The selected ICNs (i.e., network templates) were also successfully replicated by independently analyzing the data that did not pass the QC criteria, highlighting the robustness of our adaptive template to data quality.

We next studied the feasibility of estimating the corresponding subject-specific ICNs using a multivariate-spatially constrained ICA as an example of group-informed network estimation approaches. The results highlight that several factors, including ICNs themselves, data length, and spatial resolution, play key roles in successfully estimating the ICNs at the subject level. Large-scale ICNs, in general, require less data to achieve a specific level of spatial similarity with their templates (as well as within- and between-subject spatial similarity). Moreover, increasing data length can reduce an ICN’s subject-level specificity, suggesting longer scans might not always be desirable. We also show spatial smoothing can alter results, and the positive linear relationship we observed between data length and spatial smoothness (we posit that it is at least partially due to averaging over intrinsic dynamics or individual variation) indicates the importance of considering this factor in studies such as those focused on optimizing data length. Finally, the consistency in the spatial similarity between ICNs estimated using the full-length of data and subset of it across different data lengths may suggest that the lower within-subject spatial similarity in shorter data lengths is not necessarily only defined by lower reliability in ICN estimates; rather, it can also be an indication of brain dynamics (i.e., different subsets of data may reflect different ICN dynamics), and as we increase the data length, the result approaches the average (also known as static) ICN pattern, and therefore loses its distinctiveness.

## 1. Introduction

Resting-state functional MRI (rsfMRI) studies have significantly advanced our knowledge of both typical and disordered brain functional organization by evaluating the functional interactions across the brain using the blood-oxygenation-level-dependent (BOLD) signal. While rsfMRI has several advantages that make its application in a wide range of clinical and research settings more feasible than task-based fMRI paradigms, its clinical utility and application in single-subject analyses have been limited.

Clinical applications and statistical inferences are generally built upon identifying and evaluating common patterns/features. In structural MRI analysis, we investigate unambiguous brain structures where changes in the properties of a given structure can be assessed as an indication of abnormality. The task-based fMRI analysis captures the brain’s response to well-defined external tasks to identify and evaluate the task-related features across individuals. However, because the ground truth of functional entities in a given brain is unknown, the identification of the corresponding functional patterns across individuals is not straightforward for rsfMRI. The limitation of existing approaches to obtain comparable functional patterns across individuals and brain states in a way that accurately and precisely captures both individual variation and inter-subject correspondence is one primary factor limiting rsfMRI applications.

Focusing on functional connectivity, the most common category of approaches uses anatomically fixed regions (i.e., existing atlases) and evaluates the functional connectivity between these regions. By using anatomically fixed regions, this category implicitly assumes the functional (connectivity) profile within each anatomically fixed region does not vary over time and is the same across individuals. However, many static and dynamic rsfMRI studies have challenged this strong assumption by identifying differences in the spatial patterns of functional entities both across subjects and within subjects over time (Boukhdhir et al., 2021; Erhardt et al., 2011; Iraji et al., 2019a; Iraji et al., 2019c; Luo et al., 2021; Wang et al., 2015).

The presence of within- and between-subject spatial differences is further supported by task-based fMRI findings showing that functional connectivity maps and spatial patterns of brain responses vary across individuals for a given task as well as within-subject across mental states dictated by tasks (Calhoun et al., 2008; Krienen et al., 2014; Salehi et al., 2020; Sui et al., 2009; Wu et al., 2021). As a result, data-driven approaches are gaining interest because they identify functional entities from the rsfMRI data itself and therefore incorporate spatial variabilities in calculating corresponding functional connectivity patterns.

There are two main categories of data-driven approaches. They either (1) estimate functional entities for each sample (e.g., subject) and then match them across samples or (2) obtain group-level functional entities using entire samples (e.g., group-level brain networks) and then use them as a reference to estimate corresponding functional entities for each sample. Early data-driven approaches belong to the first category, including those that apply independent component analysis (ICA) to each subject’s data to extract intrinsic connectivity networks (ICNs; as estimations of functional entities) and then perform a matching step (e.g., clustering) to identify the ICN correspondence across individuals. While this category of approaches has remained a matter of great interest with significant potential (Calhoun et al., 2001a; Durieux and Wilderjans, 2019; Esposito et al., 2005; Gordon et al., 2017a; Salehi et al., 2020), ambiguity and uncertainty that the matched functional entities represent the best corresponding functional patterns across individuals remained their major drawback. Studies have shown that a slight change in the seed location could result in significant differences in functional connectivity patterns (Yeo et al., 2011), indicating that finding the best-matched patterns across individuals requires an extensive search across all possibilities. In the case of using functional parcellations as functional entities, this means searching for different sizes and locations across all individuals. In addition, extending the application to new unseen data, which is necessary for clinical application, requires special solutions as accessibility to the initial dataset and rerunning the process is not only practically unfeasible, but importantly can lead to different functional patterns than the original analysis. Differences in data acquisitions, such as different spatial and temporal resolutions, can also impact functional pattern identification. Finally, the signal-to-noise ratio (SNR) of rsfMRI adds yet another challenge, reducing the likelihood of finding the same functional entities across individuals. Finding corresponding functional entities is even more challenging when considering the dynamic nature of the brain and the fact that functional entities continuously evolve and have different spatial profiles (Iraji et al., 2020).

The second category of data-driven approaches, also known as group-informed approaches, has been deployed to overcome these limitations and enhance the identification of corresponding functional patterns across individuals (Calhoun et al., 2001b). These approaches utilize a template obtained from the data of multiple subjects to guide the estimation of corresponding functional patterns for each individual. Leveraging the data of multiple subjects provides a more reliable estimation of functional entities, and using the common template for sample estimation of functional sources improves the identification of corresponding patterns across individuals. As a result, this category can be more beneficial for widespread clinical adoption as it enhances the possibility of comparing the same functional patterns (and their dynamic states) across individuals. Group ICA + back-reconstruction is the most commonly used example of this category, which uses group-level ICNs estimates to obtain corresponding subject-level ICNs (Erhardt et al., 2011). Yet, two aspects must be improved to fully leverage this category’s potential.

First, we need to obtain a reliable template (e.g., group-level ICNs) that best represents all individuals. A key factor in achieving this goal is to recruit the largest possible dataset. As we increase the size of a dataset, the group-level estimations get closer to the central tendency, and therefore, better represent all (seen and unseen) individuals. Second, we need to use well-designed group-informed (e.g., reference-guided or back-reconstruction) network estimation techniques to identify corresponding subject-specific functional patterns accurately and precisely. This step is important to prevent loss of subject-specificity and meaningful inter- (and intra-) individual differences. Multiple studies have shown that existing techniques capture individual-level variations well (Allen et al., 2012; Erhardt et al., 2011), yet developing more advanced network estimation techniques (in addition to accurate estimation of a template) is a key element to bringing group-informed approaches to perfection and transitioning to clinical applications of rsfMRI.

In other words, a standardized framework that leverages a very large dataset to obtain a reliable general template of functional entities and uses techniques that allows accurate subject-specific estimation of these functional entities can lead to the systematic characterization of common and distinct alterations in functional patterns across cohorts (including among clinically overlapping disorders) and identifications of subject-specific irregularities. This standardized framework makes identifying corresponding functional patterns for new subjects and comparing findings among datasets and across studies straightforward, which is of great need in the field. Moreover, because the estimation for each subject is independent of other subjects in the study, it becomes an ideal solution for both clinical applications as well as prediction analysis, which requires complete separation of the training and testing data.

This study focuses on fulfilling the first piece of this standardized framework, i.e., identifying a reliable global ICN template. Towards this goal, we use a large dataset (over 100k subjects) and group multi-model-order ICA to generate a set of common, reliable multi-spatial-scale ICNs. Compared to previous similar attempts (Du et al., 2020), our work uses a much larger sample size, obtains ICNs across multiple spatial scales, provides more reliable and replicable ICNs, and does not restrict to typical control cohorts. We chose to use all data available to us (including clinical cohorts) to ensure the obtained ICN template reflects the diversity and heterogeneity of the brain and can be broadly representative of different groups. In this framework, obtaining a universal template that best represents all individuals is the primary objective for the template, and estimating accurate subject-specific networks is the primary objective of the group-informed network estimation technique. We evaluate the identification of the corresponding subject-specific ICNs using multivariate-objective optimization ICA with reference (MOO-ICAR) (Du and Fan, 2013) and present basic tools and metrics to test the successful identification of subject-specific ICNs for a given dataset. Applying group-informed approaches to a given dataset does not guarantee the successful identification of corresponding functional patterns. There is a significant need to evaluate the factors that influence sample-specific estimates and also provide tools for future applications. In this, we took our first step toward this important realm of research.

## 2. Materials and Methods

### 2.1. Datasets and data preparation

#### 2.1.1. Dataset

We utilized rsfMRI data from 100,517 subjects available in more than twenty private and public datasets. The full list of datasets and resources for obtaining further details on each can be found in Supplementary 1. Datasets are from cohorts with different male-to-female ratios, age distributions, handedness, and diagnosis, collected by different scanners with varying imaging protocols such as different spatial and temporal resolutions. In this initial work, we leveraged as much data as possible to identify an ICN template, and the influence of various demographic factors will be evaluated in future studies, given the demographic distribution availability. Here, we focused on data quality control (QC) criteria to screen and select data without further exclusion criteria. The QC criteria include (a) a minimum of 120 time points (volumes) in the rsfMRI time series, (b) mean framewise displacement less than 0.25, (c) head motion transition less than 3° rotation and 3 mm translation in any direction, (d) high-quality registration to an echo-planar imaging template, and (e) whole-brain (and the top ten and the bottom ten slices) spatial overlap between the individual mask and group mask above 80%. We chose these QC criteria as they are both reasonable and achievable across different datasets. This resulted in 57,709 (57.4%) individuals who passed the QC criteria, which we called the QC-passed dataset, in contrast to the QC-failed dataset defining the remaining 42,808 (42.6%) individuals who did not pass the QC criteria. We used the QC-passed dataset to extract the ICN template and evaluated the replicability and presence of selected ICNs by separately analyzing the QC-failed dataset.

#### 2.1.2. Preprocessing

If the preprocessed data were available for a given dataset, we used the preprocessed data; otherwise, we performed preprocessing steps, including rigid body motion correction, slice timing correction, and distortion correction, using a combination of FMRIB Software Library (FSL v6.0, https://fsl.fmrib.ox.ac.uk/fsl/fslwiki/) and the statistical parametric mapping (SPM12, http://www.fil.ion.ucl.ac.uk/spm/) toolboxes under the MATLAB environment. Next, preprocessed subject data were warped into a Montreal Neurological Institute (MNI) space using an echo-planar imaging (EPI) template, as it has been shown to outperform structural templates (Calhoun et al., 2017) when distortion correction is unavailable or unfeasible, as was the case for this study. Finally, subject data were resampled to 3 mm^3^ isotropic voxels and spatially smoothed using a Gaussian kernel with a 6 mm full width at half-maximum (FWHM).

### 2.2. Intrinsic Connectivity Network (ICNs) Template Estimation

The analysis pipeline is displayed in Figure 1. Using our QC-passed dataset, we applied group-level multi-model-order spatial ICA (gr-msICA) (see section 2.2.1.) to obtain a multi-model-order ICN template. For this purpose, we first randomly half split the QC-passed data and applied gr-msICA on each half independently. We used model orders of 25, 50, 75, 100, 125, 150, 175, and 200, totaling 900 independent components (ICs) for each data split. We repeated this process 50 times, generating 100 sets of 900 ICs. Next, we applied a greedy search and selected the 900 ICs with the highest average pairwise spatial similarity (calculated by Pearson correlation) across 100 sets. It is worth mentioning that using more granular model order increments and including higher ICA model orders (Iraji et al., 2019b; Smith et al., 2013) is advantageous to improving the reliability of ICN estimations and obtaining a more complete set of reliable ICNs across a broader range of spatial scales. However, implementing these two elements is computationally very expensive and becomes impractical considering the large number of ICA runs in our proposed stability analysis pipeline. Finally, we manually labeled the ICs and selected 105 of the ICNs that are different from each other with all had spatial similarity below 0.8 with each other to serve as the final ICN template (see section 2.2.2.).

**Figure 1.**
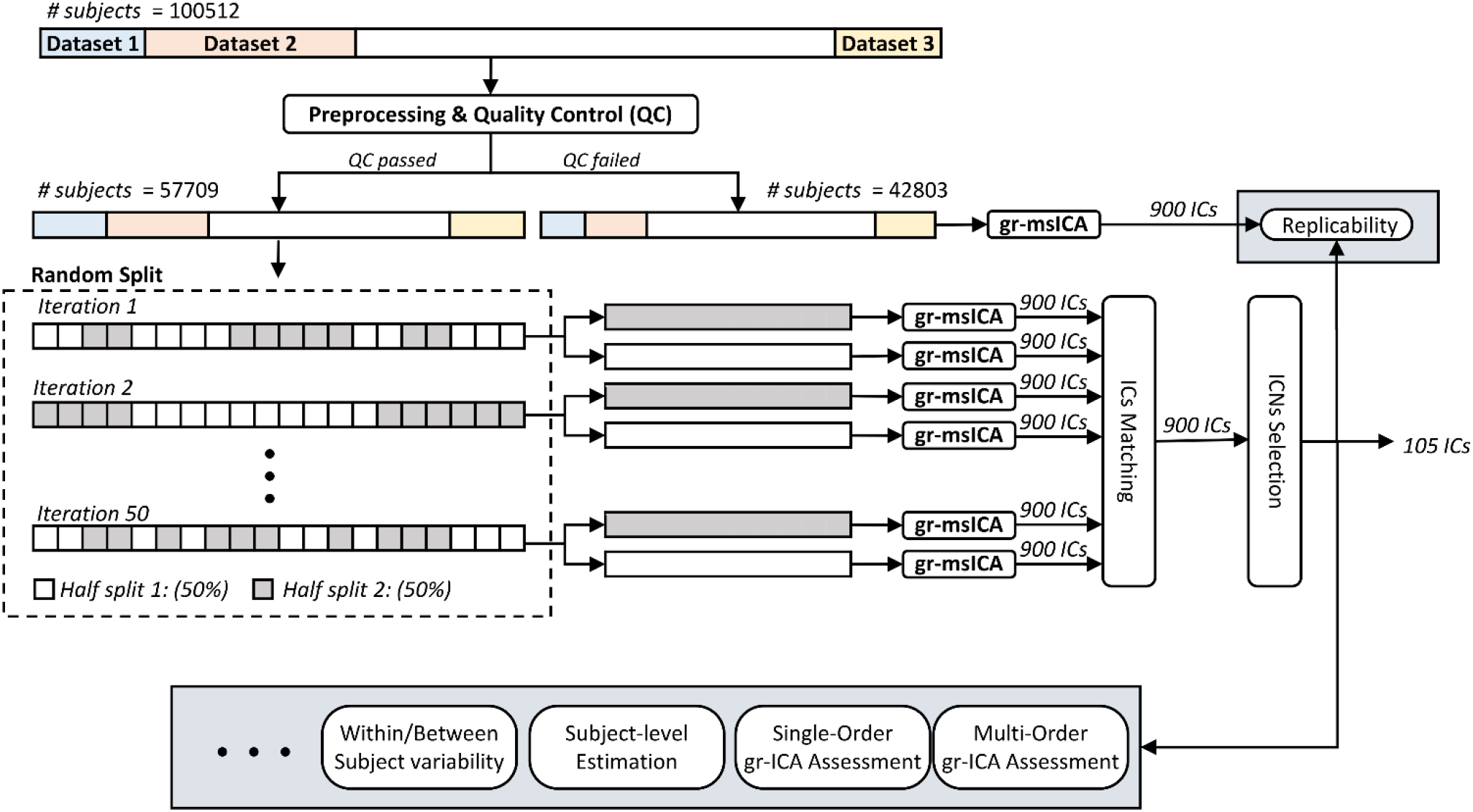
Analysis pipeline. Data of 100,512 subjects went through preprocessing and quality control (QC). In total, 57,709 subjects passed the QC and were used to generate the template. The QC-passed dataset was randomly split in half, and group-level multi-spatial-scale independent component analysis (gr-msICA) was applied on each half split to generate 900 independent components (ICs). This process was repeated independently 50 times, which resulted in 100 sets of 900 ICs. Next, the 900 most stable ICs were identified and labeled as non-ICNs or ICNs, and the 105 most distinct (spatial similarity < 0.8) were selected as the ICN template. Finally, several group-level and subject-level analyses were performed.

#### 2.2.1. Group-level Multi-Model-Order Spatial Independent Component Analysis (gr-msICA)

ICA analysis was performed using the Group ICA of FMRI Toolbox (GIFT) v4.0c package (https://trendscenter.org/software/gift/) (Iraji et al., 2021). Spatial ICA is a multivariate blind source separation technique that simultaneously considers the relationships among all voxels (as opposed to pairwise Pearson correlations) to estimate temporally coherent spatial patterns that are maximally independent for a selected model order. The group ICA analysis steps are as follows. We first applied variance normalization (z-score) on voxel time courses and computed subject spatial principal components analysis (PCA) to retain the principal components (PCs) with maximum subject-level variance (greater than 95%). Next, group spatial PCA was applied to stacked subject PCs to obtain subject commonalities and subspace with the maximum variation across the whole dataset. Group PCs were computed using a memory-efficient subsampled time PCA (STP) approach (Rachakonda et al., 2016). We used STP because the conventional group spatial PCA is intractable considering the data size used in this study. STP estimates the group PC subspace by incremental updating based on a different sub-stack of subject PCs. In other words, first-level group PCA was applied to different subsets of subject PCs, and then the final group PC subspace was estimated by incrementally updating and incorporating first-level group PCs (Rachakonda et al., 2016). Next, we ran gr-msICA using the Infomax ICA algorithm (Bell and Sejnowski, 1995) with model orders of 25, 50, 75, 100, 125, 150, 175, and 200. We ran Infomax 20 times for each model order to obtain the most stable run for each model order (Du et al., 2014).

We used msICA to extract ICNs that exist across different spatial scales (Iraji et al., 2022), from large-scale spatially distributed ICNs (Damoiseaux et al., 2008; Iraji et al., 2016) to more spatially granular ICNs (Allen et al., 2011; Iraji et al., 2019b). It is worth noting that the model order of ICA effectively sets the spatial scale of ICNs without imposing a direct spatial constraint (Iraji et al., 2022). This is a great advantage as the complexity varies across brain systems, and there is no reason to expect distinct regions (e.g., temporal lobe vs. the frontal lobe) or systems (e.g., the visual vs. the cognitive control) to have the same spatial scale across the brain functional hierarchy.

The second advantage of using msICA is the superior ability to estimate more reliable ICNs across different datasets. Briefly, the amount of variance that can be explained by a given IC relative to other ICs can vary across datasets, which impacts the PCs retrieved by group-level PCA steps, and therefore the input for ICA decomposition. In other words, subject variabilities influence the data reduction steps prior to ICA decomposition, and hence the output of ICA. Variability in ICA estimations, such as statistical errors and several equally good local minima solutions, also impact the output of ICA decompositions, which we commonly minimize by running ICA several times and identifying the best run (Ma et al., 2011). Nonetheless, these sources of variability together lead to a better estimation of a given ICN at different model orders across different datasets. In layman’s terms, a given ICN can be best identified either in one specific model order across datasets or in different model orders. Thus, by using ICA with multiple model orders, we can improve the identification of ICNs across datasets. However, due to computational overhead, we only use a limited number of model orders with a relatively large step size of 25 (i.e., 25, 50, 75, 100, 125, 150, 175, and 200), so we can only leverage this advantage of the multi-model-order framework as far as computational feasibility allows.

#### 2.2.2. ICN Selection

We applied msICA on 50 random half-splits of QC-passed data and obtained 100 sets of 900 ICs. Next, we identified the most stable ICs across the 100 runs. ICASSO is the most commonly used tool to find stable ICs and its extension for multi-model-order ICA applications is straightforward. However, the computational complexity of ICASSO rapidly increases with the number of ICs and ICA runs, making it impractical for our study (900 × 100 IC samples). As such, we applied a procedure that selects ICs based on the best pairwise matching of ICs between runs. The steps of the procedure are as follows. First, for each IC of each run, we found its best-matched component from all the other 99 runs. A best-matched component was defined as the component with maximum spatial Pearson correlation. We calculated the average of the 99 correlation values of best-matched components as the stability index for that given component. We obtained this stability index for all components across all runs resulting in a 900 × 100 stability matrix. Next, we identified the component with maximum stability value across all 900 × 100 components as the first selected component, and the selected component along with its best-matched component in each run were removed from search space (i.e., for each run, the component with maximum similarity with the selected component will be identified and omitted for the next iteration). We repeated the whole procedure for the remaining 899 × 100 components and continued this procedure until we selected 900 components along with their corresponding components across all runs. Next, four authors (V.C., Z.F., A.F., and A.I.) manually labeled the 900 selected components as ICNs or non-ICNs.

Note that some ICNs might have high spatial similarities with each other, and thus we chose a subset of ICNs (*N* = 105) with spatial similarity less than 0.8 as the ICN template (Supplementary 2). All 900 components and their stability values can be found in Supplementary 3 to allow researchers to select criteria that best match their objectives.

### 2.3. Subject-level Estimation of ICNs

Several factors play roles in estimating corresponding ICNs at the subject level, including fMRI data characteristics, ICNs themselves, and group-informed network estimation techniques. The data characteristics, such as the amount (i.e., number of time points) of subject-level data and spatial as well as the temporal resolutions of data, define the limits of ICNs estimation. In other words, a given ICN cannot be estimated if enough information is not present in the data, regardless of group-informed network estimation techniques. The properties of each ICN (e.g., its spatial distribution, the amount of data variance it explains, and how densely its main cores are temporally coupled) determine how easily they can be extracted. In other words, the spatial and temporal information required to identify each ICN varies. Some ICNs can be estimated using fewer time points and coarse spatial and temporal data, while others may require fine-grained spatial and temporal information. Group-informed network estimation techniques are the determinant factor in obtaining the best estimation of subject-level ICNs for a given fMRI time series. As such, there is a significant effort to develop robust, reliable methods with a high level of sensitivity and specificity to simultaneously and accurately identify corresponding ICNs for a given sample data while capturing sample-specific (e.g., subject-specific) fine information (Du and Fan, 2013; Lin et al., 2010; Mejia et al., 2020). While we emphasize the necessity of developing new, more advanced methods, we leave this effort to future endeavors and use MOO-ICAR to estimate subject-specific ICNs because it is suggested to perform well in capturing subject-specific information and removing artifacts (Du et al., 2016; Du and Fan, 2013).

All these factors together highlight that merely using an ICN template and a group-informed network estimation technique does not guarantee proper estimation of subject-level ICNs, and therefore underscores the need to use criteria to evaluate the success of the ICN estimations. Here, we proposed two minimum criteria to systemically evaluate how effectively a given ICN is estimated in a given dataset. The first criterion evaluates whether the spatial similarity between a given ICN’s template and its subject-level estimates in a given dataset is significantly higher than the spatial similarity between the ICN’s template and components estimated from null data with the same data length and level of spatial smoothing. This criterion basically determines whether an ICN is estimated significantly beyond just using predefined anatomical information determined by its template. Otherwise, the estimations reduce to using predefined spatially fixed weighted nodes/seeds (i.e., become equal to atlas-based approaches). The second criterion evaluates if an estimated ICN has significantly higher spatial similarity to its own template compared to its similarity to templates from other ICNs. Data specifications (e.g., resolutions and length) and group-informed network estimation techniques determine the ability to differentiate between ICNs.

We evaluate the success of estimating ICNs using the two introduced criteria for the Functional Imaging Biomedical Informatics Research Network (FBIRN) (Keator et al., 2016) and Human Connectome Project (HCP)(Van Essen et al., 2013) datasets with different specifications (including inherent spatial resolution). The FBIRN dataset contains 109 subjects that passed QC with 162 time points, a repetition time (TR) of 2 sec, and an original voxel size of 3.4375 × 3.4375 × 4 mm^3^. The HCP dataset consists of 706 subjects that passed QC and have four complete rsfMRI scan sessions. Each session has 1200 time points, a TR = 0.72 sec, and an original voxel size of 2.0 mm isotropic. We also investigated ICN estimation in the context of (a) similarity to template ICNs, (b) within-subject similarity, and (c) between-subject similarity for different data lengths and sampling rates. We focused on the HCP dataset for data length and temporal resolution assessments. We discarded the first 50 time points and partitioned the data from each session into incrementally longer segments, beginning at 25 time points with an increment of 25 time points (i.e., 25, 50, 75,…, 1150). We performed this process for data with different temporal resolutions of 1 to 5 TR (0.72, 1.44,…, 3.6 sec). We applied MOO-ICAR separately to each data length and temporal resolution to estimate the corresponding 105 ICNs.

### 2.4. On smoothing and reliability analysis

Spatial smoothness induced by sample size is an understudied research area. For instance, the group ICA results, similar to averaging and any group-level analysis, are smoother than the single-subject ICA results, as sample-specific spatial variation is gradually smoothed out as more data are included. This phenomenon can also be observed when calculating the functional connectivity map of a given seed and averaging it across individuals. This smoothing effect can also be seen when increasing the sample size (number of time points in this case) for estimating the subject-level functional connectivity (FC) patterns. The more time points that are included, the smoother the FC patterns (e.g., spatially more smoothed ICNs) become.

This smoothness associated with data lengths directly impacts the results of downstream assessments (e.g., reliability assessment) that use spatial similarity to evaluate a system’s performance, since more spatially smoothed estimations in general result in higher spatial similarity. Therefore, the impact of this smoothing effect should be factored in for reliability analysis and similarity evaluations, especially when comparing results from data with different data lengths. Otherwise, the results are always biased toward longer data lengths and therefore support the notion of using longer data lengths. A fair comparison between the results of different data lengths may need to control for the induced spatial smoothness due to this averaging effect. One straightforward solution is to match the smoothness of ICN estimations across different data lengths, for instance, by applying different levels of spatial smoothing for different data lengths to achieve a similar level of overall smoothing across data lengths.

At the same time, one should take caution about the extent to which the spatial variations are driven by sample variability and error of estimation, or to what extent they reflect a true underlying pattern. While higher spatial similarity among a given ICN’s estimations (such as similarity to its reference and its within-subject and between-subject similarity) is an appealing feature for many analyses, for example, when using spatial similarity as reliability or replicability criteria, it could potentially come at the cost of reducing subject differences, fine spatial information, and dynamic properties. Thus, applying spatial smoothing (to match smoothness across analyses) in studies could also include an assessment of its impact not only on reliability and replicability analysis but also its impact on other analyses such as prediction and association. Given the extensive results already included in this paper, we plan to incorporate a rigorous analysis on the effects of spatial smoothing equalization across different data lengths for future studies. Nonetheless, in the present work, we conducted some baseline analyses to demonstrate the general impact of spatial smoothing on subject-level ICN evaluations. For this purpose, we applied *post hoc* smoothing using a Gaussian kernel with a fixed full width half maximum (FWHM) to the subject-level ICNs estimated across all data lengths, i.e., the same level of smoothing (FWHM = 4.9 mm) was applied to all ICNs, regardless of the data length used for estimation. The FWHM value was selected as the value that gave the highest average spatial similarity to the template ICNs across all data lengths in independent external data. We evaluated how smoothing affects the results of our analyses, such as the similarity between subject-level ICNs and their template, as well as both within-subject and between-subject similarities.

## 3. Results

### 3.1. Reliable Intrinsic Connectivity Network (ICNs) Template

Figure 2(A) displays the composite views of the 105 selected ICNs and their average functional connectivity. The stability index (average spatial similarity across runs) for all ICNs is well above 0.8. Figure 2(B) shows the spatial similarity between ICNs and best-matched components across 100 runs of gr-msICA on 50% of the QC-passed dataset. We additionally ran gr-msICA on the QC-failed dataset, which was not used in ICNs estimation (unseen dataset), and successfully (spatial similarity > 0.8) identified all 105 ICNs.

**Figure 2.**
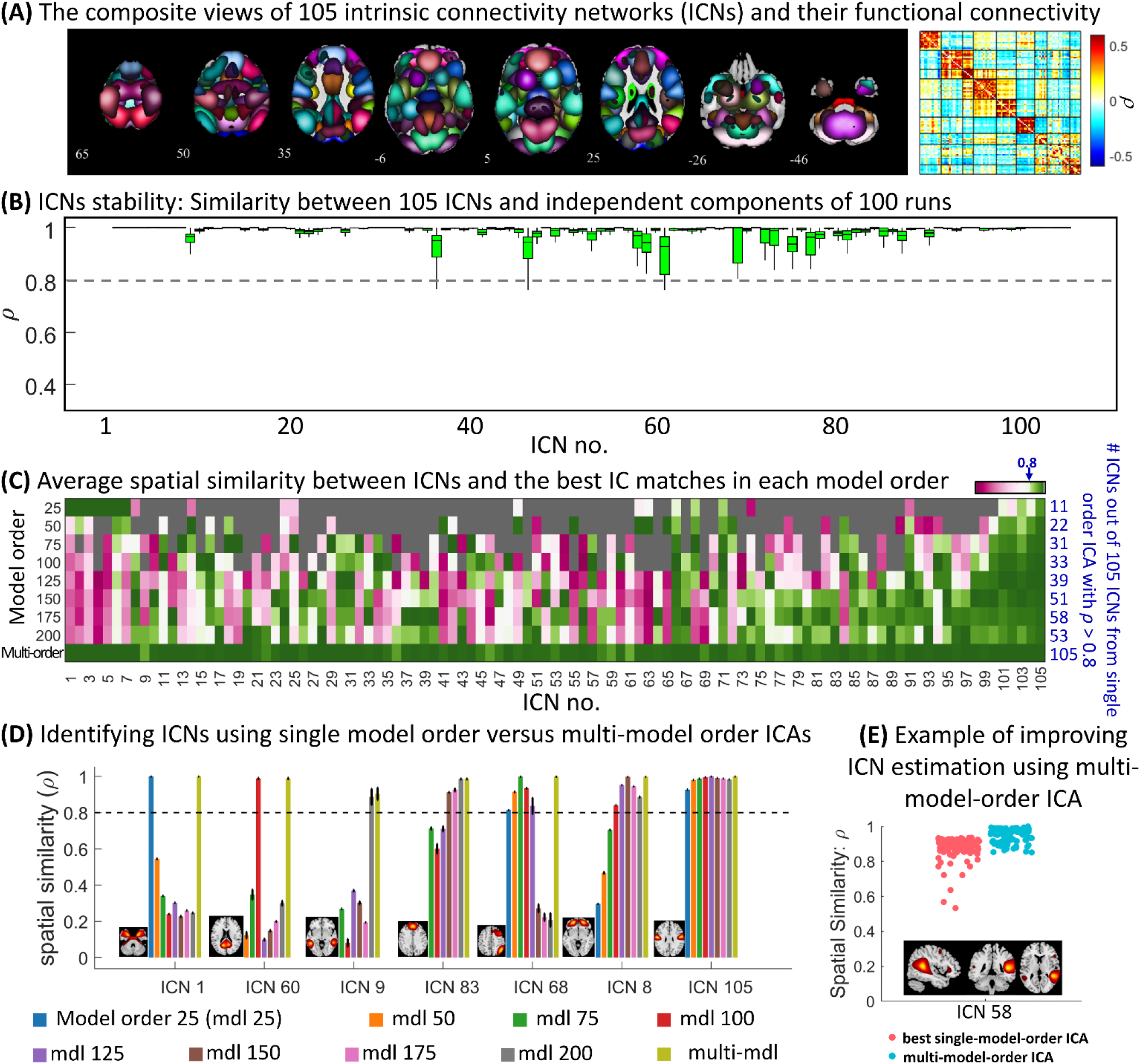
Group-level multi-spatial-scale independent component analysis (gr-msICA) results. (A) The composite views of the 105 selected intrinsic connectivity networks (ICNs) and averaged whole-brain functional connectivity. Each ICN spatial map was first z-scored and thresholded at z-value = 1.96 (p-value = 0.05). Whole brain functional connectivity was estimated by calculating the Pearson correlation between each pair of ICNs and averaged across the QC-passed dataset. (B) Multi-model order stability. The spatial similarity of ICNs with corresponding independent components (IC) across 100 gr-msICA runs on different halves of the QC-passed dataset. (C) single-model-order versus multi-model-order ICA. The average spatial similarity is computed between each ICN and the best IC matches across the 100 runs in each model order (and multi-model-order). The blue values on the right side indicate the number of ICNs identified by each model order using the similarity threshold of 0.8 (similar to stability index = 0.8). (D) Different ICNs can be identified using different ranges of model orders. For example, ICN 105 was successfully identified by all model orders used in this study. (E) an example showing how using msICA improves the identification of the best corresponding ICN across different subsets of data, suggesting msICA improves the identification of ICNs across datasets.

Our results suggest that leveraging the msICA framework can improve ICN identification. Figure 2(C) shows the average spatial similarity between each ICN and the best IC matches across 100 runs in each model order as well as the number of ICNs identified by each model order using the similarity threshold of 0.8 (similar to stability index = 0.8). This analysis shows that (1) no single model order ICA can estimate all of the 105 ICNs (e.g., only 11 out of 105 ICNs were identified using model order 25 and only about half (58) of the ICNs were identified using model order 175); therefore, msICA provides a more complete view of brain functional patterns and (2) the ICNs represent different patterns across the various ICA model orders, meaning each can be identified from a different range of model orders. Examples of these differences can also be seen in Figure 2(D). For example, ICN 105 was successfully identified across all model orders, while ICNs 1, 9, and 60 were only identified in one model order. We also observed that some ICNs were successfully identified (ρ ≥ 0.8) across multiple model orders, with one model order exhibiting the highest stability value (e.g., ICN 8 and 68).

Figure 2(E) provides an example of a single ICN where the best-matched components across runs (which contain different data) come from different model orders. In this example, we can obtain ICN 58 using a single-model-order ICA with a stability index above 0.8; however, because the best-matched components can come from different model orders, we achieved higher stability and improved the identification of ICNs across datasets by leveraging multi-model-order ICA, which is the second abovementioned advantage of multi-model-order ICA. Our analysis shows that 28 of the 105 ICNs came from different model orders (2 to 4 different model orders) across 100 half-split runs. We further investigated this benefit of multi-model-order ICA by comparing the results of applying msICA on the QC-failed dataset with 100 runs from the QC-passed dataset. Our findings showed that for 53 out of the 105 ICNs, the corresponding ICNs in the QC-failed dataset were identified in different model orders than all QC-passed runs.

### 3.2. Subject-Level Analysis

The subject-level analysis suggests that we can identify subject-specific ICNs corresponding to our template using existing data acquisition paradigms and back-reconstruction methods. MOO-ICAR estimated all ICNs above null for all HCP data lengths (i.e., 25, 50, 75,…, 1150 time points); however, it could not effectively extract ICN-specific information for 6, 5, and 2 ICNs for the shortest data lengths of 25, 50, and 75 time points, respectively, meaning the MOO-ICAR solutions for these ICNs were not statistically (p-value > 0.05) more spatially similar to their templates compared to that of other templates.

Indeed, the results of subject-level estimations can vary based on both back-reconstruction and data characteristics. For instance, for the FBIRN dataset with 157 time points and a larger original voxel size of 3.4375 × 3.4375 × 4 mm (i.e., higher inherent spatial smoothness), MOO-ICAR estimation of 12 ICNs did not show statistically higher similarity to their reference than other references. In comparison, for the HCP dataset with data length equal to or larger than 100 time points, MOO-ICAR successfully estimated all 105 ICNs.

In Figure 3(A), the top row shows an example of subject-level ICN estimation from both the HCP and FBIRN datasets. Subject-specific estimations have less spatial smoothness than the ICN template obtained from the group-level analysis. Subject-specific smoothness gradually increases as more time points are used for MOO-ICAR estimations (Figure 3(B)). In other words, we illustrated the expected increase phenomena (see section 2.4) in spatial smoothness (computed as one minus average gradient magnitude across the whole brain) as a function of data length for an exemplar ICN. We also employed the post hoc subject-level spatial smoothing (see section 2.4) and observed that this smoothing indeed increased the similarity between subject-level ICNs and templates obtained from group-level analysis (the bottom row of Figure 3(A) provides a visual illustration).

**Figure 3.**
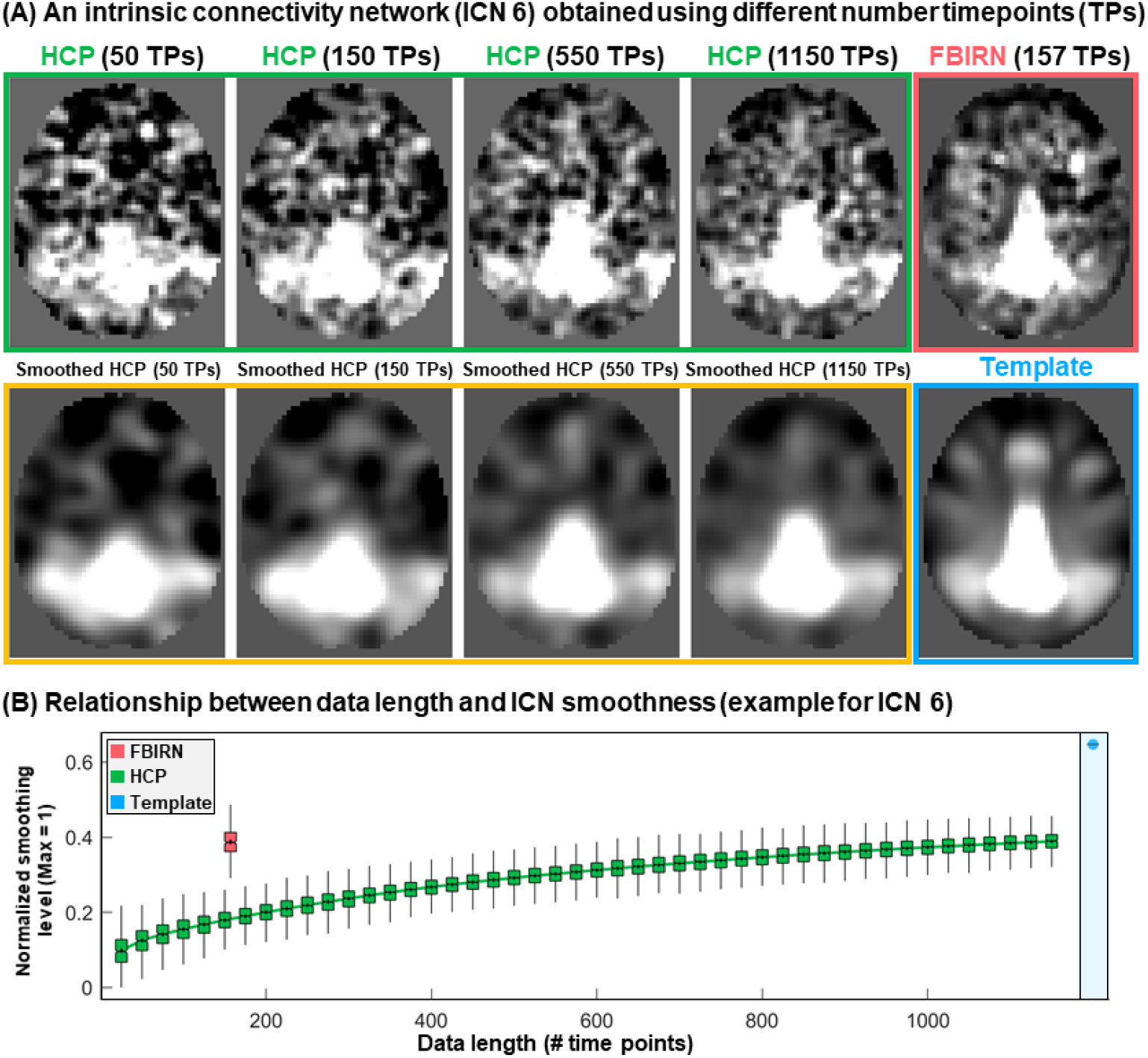
Subject-level estimation and the effect of smoothness. (A) Estimation of intrinsic connectivity network 6 (ICN 6) for a single subject from the Human Connectome Project (HCP) dataset using different data lengths and one subject from the Functional Imaging Biomedical Informatics Research Network (FBIRN). The subject-level estimation is less smooth than the template obtained using group-level analysis. The subject-level estimate depends on several factors, including data characteristics such as length of data and original voxel size. (B) The smoothness of ICN 6 estimated as a function of data lengths for the HCP dataset is shown in green. The red color represents the same measure for the FBIRN dataset with 157 time points. Blue shows the smoothness level for the template of ICN 6. Y-axis shows the normalized smoothness level with a maximum value of 1, which corresponds to a constant image (when all voxels have the same value, the normalized smoothness level is equal to one). Normalized smoothness equals one minus average gradient magnitude across the whole brain.

The results of the spatial similarity analysis between the subject-specific ICNs and the template are summarized in Figure 4. The left column shows the results for the original ICN estimations, and the right column represents assessments after applying the post hoc spatial smoothing (i.e., applying the same spatial smoothing with FWHM = 4.9 mm to all ICNs and data lengths). The first row (Figure 4(A)) shows the average similarity of 105 ICNs with the template, where the green shaded area represents its standard deviations across individuals, and Figure 4(B) shows an example of the spatial similarity for ICN 38. The pattern of spatial similarity with the template varies across ICNs. Figure 4(C) shows differences in the similarity of selected ICNs with the template, indicating different ICNs need different data lengths (i.e., numbers of samples) to achieve a specific spatial similarity to the template and spatial smoothing increases the similarity across all ICNs.

**Figure 4.**
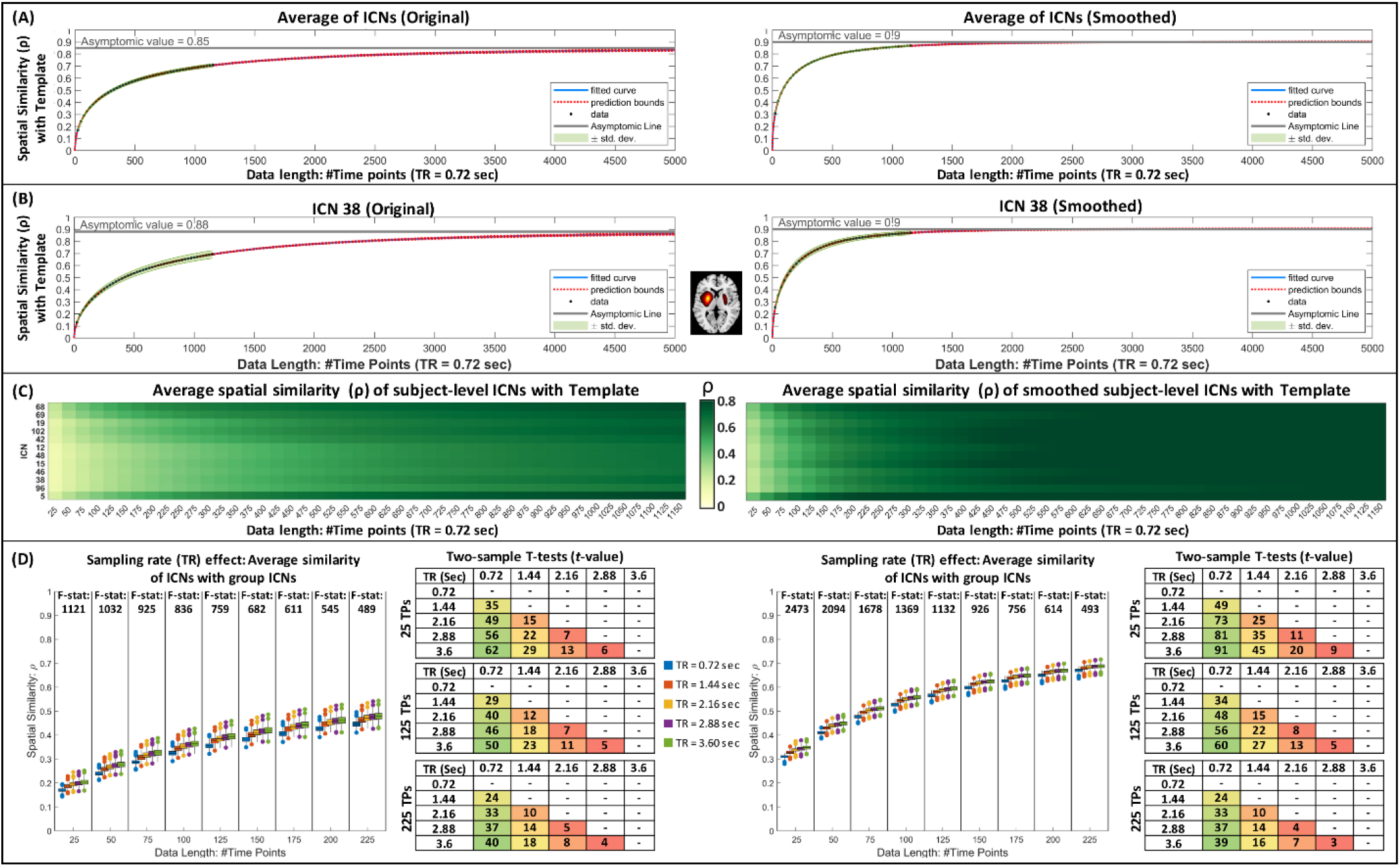
Assessment of subject-level intrinsic connectivity networks (ICNs) estimated using multivariate-objective optimization ICA with reference (MOO-ICAR). The left and right columns are the results of original ICNs estimations and after applying post hoc spatial smoothing. (A) The average spatial similarity between subject-level estimations and the template using different data lengths (from 25 to 1150 time points). The green shaded area represents its standard deviations, and the blue and red dot lines represent the fitted curve and its extrapolation. (B) The same plot as (A) but for one ICN as an example. (C) While the spatial similarity between single-subject estimations and templates increases as the length of data increases, different ICNs show different patterns. Here, we show an average similarity as a function of time for a few randomly selected ICNs with and without spatial smoothing. (D) The sampling rate (TR) effect on the spatial similarity between template and subject-level estimations.

We also evaluated the effect of the sampling rate (Figure 4(D)). For the same number of time points (i.e., the same number of samples or the same data length), lower sampling rates (i.e., a longer amount of time between samples) result in slightly (but statistically significant) higher spatial similarities with the template. The effect of sampling rate is more significant in smaller data lengths. For example, the F-statistic between data with TR = 0.72, 1.44, 2.16, and 3.6 sec was 1121 for the data length of 25 time points compared to 489 for the data length of 250 time points. Furthermore, for the evaluated TR values, the two-sample t-test indicates that the impact of sampling rate reduces as sampling rate (TR) increases. For example, for the original estimation with 25 points, the t-value between the sampling rates of 0.72 and 1.44 seconds is 35, while this value is 22 for the sampling rates of 1.44 and 2.88 seconds. However, the increase in spatial similarity as a function of sampling rate seems trivial, especially considering we can (1) collect more data for the same amount of time using shorter TR and (2) better assess the changes in functional patterns over time.

Within-subject and between-subject analyses were consistent with the results of the similarity analysis with the template. For instance, we calculated the within-subject spatial similarity between different sessions of the HCP data and observed an increase in the within-subject spatial similarity as a function of data length (Figure 5(A)). Furthermore, post hoc spatial smoothing enhanced within-subject and between-subject spatial similarities (Figure 5(B)).

**Figure 5.**
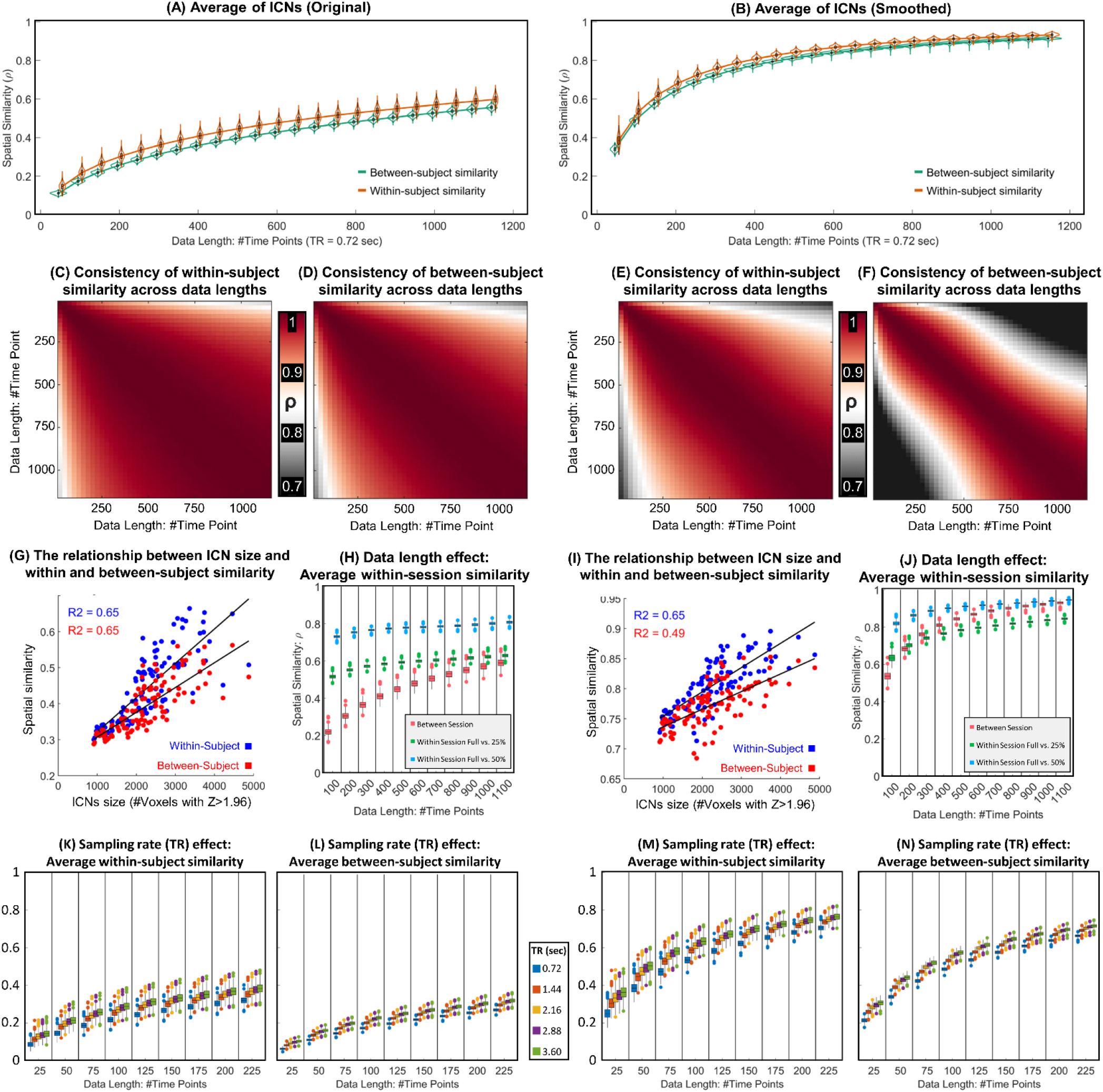
Assessment of within- and between-subject similarities. The left and right columns show the results of original ICNs estimations and after applying spatial smoothing. (A) and (B) the average within- and between-subject spatial similarity using different data lengths (from 25 to 1150 time points). (C) and (E) show the similarity of the ICNs’ within-subject spatial similarity across different data lengths for original and post hoc smoothed ICNs. (D) and (F) show similar results for between-subject spatial similarity. These results suggest that while within- and between-subject similarity is positively correlated with the data lengths (i.e., higher similarity with longer data lengths), the pattern of similarity across ICNs is consistent for different data lengths. (G) and (I) show a strong positive correlation between ICNs size and within-as well as between-subject similarity. (H) and (J) show the impact of data length on the similarity of ICNs obtained using full data lengths with those obtained using a portion (25% or 50%) of data. While the within-session similarity increases as a function of data length, this increase is not substantial, particularly compared to between-session similarity. (K) to (N) demonstrate the impact of sampling rate on within- and between-subject similarities for original analysis as well as post hoc smoothed ICNs.

While increasing data length increases within- and between-subject spatial similarities, this pattern of within- (and between-) subject similarity across ICNs remains fairly consistent across data lengths, with a higher consistency for within-subject analysis. In Figure 5(C), we calculated the average within-subject spatial similarity for all 105 ICNs at each data length. Then, we measured the Pearson correlation between 105 similarity values for each pair of data lengths. Figure 5(D) shows the same analysis for between-subject analysis, and Figure 5(E) and (F) illustrate the same analyses for post hoc spatially smoothed data. We also observed that within- and between-subject similarities are positively correlated with the size of ICNs, meaning within- and between-subject similarities seem to be higher for larger-scale ICNs than spatially more granular ICNs. Figure 5(G) and (I) illustrate the relationship between the number of voxels with Z > 1.96 (p-value < 0.05) and within- and between-subject spatial similarities.

We also evaluated the impact of data lengths and smoothing in within-session similarities relative to between-session similarities (Figure 5(H) and (J)). For this purpose, we calculated the spatial similarity of the ICNs estimated using the given data length with those estimated using a portion (25% and 50%) of that data length. For example, for the data length of 100 time points, we calculated the similarity between ICNs estimated using 100 time points with ones estimated using the initial 25 and 50 time points. Within-session similarities are more similar across data lengths, compared to between-session analysis, particularly for original estimations (Figure 5(H)). For instance, the Pearson correlation between full-length and 50% remains around 0.8 across different data lengths. Moreover, post hoc spatial smoothing has a larger impact on between-session similarities than within-session similarities. Our analysis shows the increase in spatial similarity as the result of spatial smoothing is more prominent in between-session similarities compared to within-session similarities.

The effect of the sampling rate within and between-subject similarity (Figure 5(K), (L), (M), and (N)) is also similar to what we observed in the context of similarity to the template. For the same number of samples, lower sampling rates (i.e., a longer amount of time between samples) result in overall higher within- and between-subject spatial similarities. However, the ability to collect more data for the same amount of time favors using higher sampling rates.

For the last analysis, we evaluated subject-specificity by comparing within- and between-subject similarities (Figure 6). We observed that ICNs show different patterns and levels of subject specificity. For example, Figure 6(A) shows a few ICNs with different maximum *t*-values and different relationships between data length and ICN’s subject specificity power. The maximum *t*-value is an indication of a given ICN’s ability to differentiate between individuals, and we observed, for example, this value is much higher for ICN 68 (commonly known to belong to the frontoparietal domain) than ICN 5, which is a large-scale ICN in the cerebellum. Moreover, the *t*-value of within- and between-subject similarity gradually increases as a function of data length for ICN 68 in explored data length, while the value reaches the maximum in mid-range data lengths for ICN 5. Figure 6(B) shows that the pattern of ICNs’ subject-specificity remains fairly similar across different data lengths, particularly when the data length is above 500 time points. Similar to the previous analyses, subject-specificity is positively correlated with the ICN spatial extent, meaning larger scale ICNs estimated using MOO-ICAR carry more subject-specificity power relative to spatially finer scale ICNs (Figure 6(C)). Further investigation shows that subject-specificity variability was non-uniformly distributed across systems. ICNs associated with frontoparietal and default mode show higher within-versus-between subject *t*-values, while ICNs associated with the subcortical and cerebellum have the lowest subject-specificity power, and this pattern is similar for the original as well as post hoc smoothing (Figure 6(D)).

**Figure 6.**
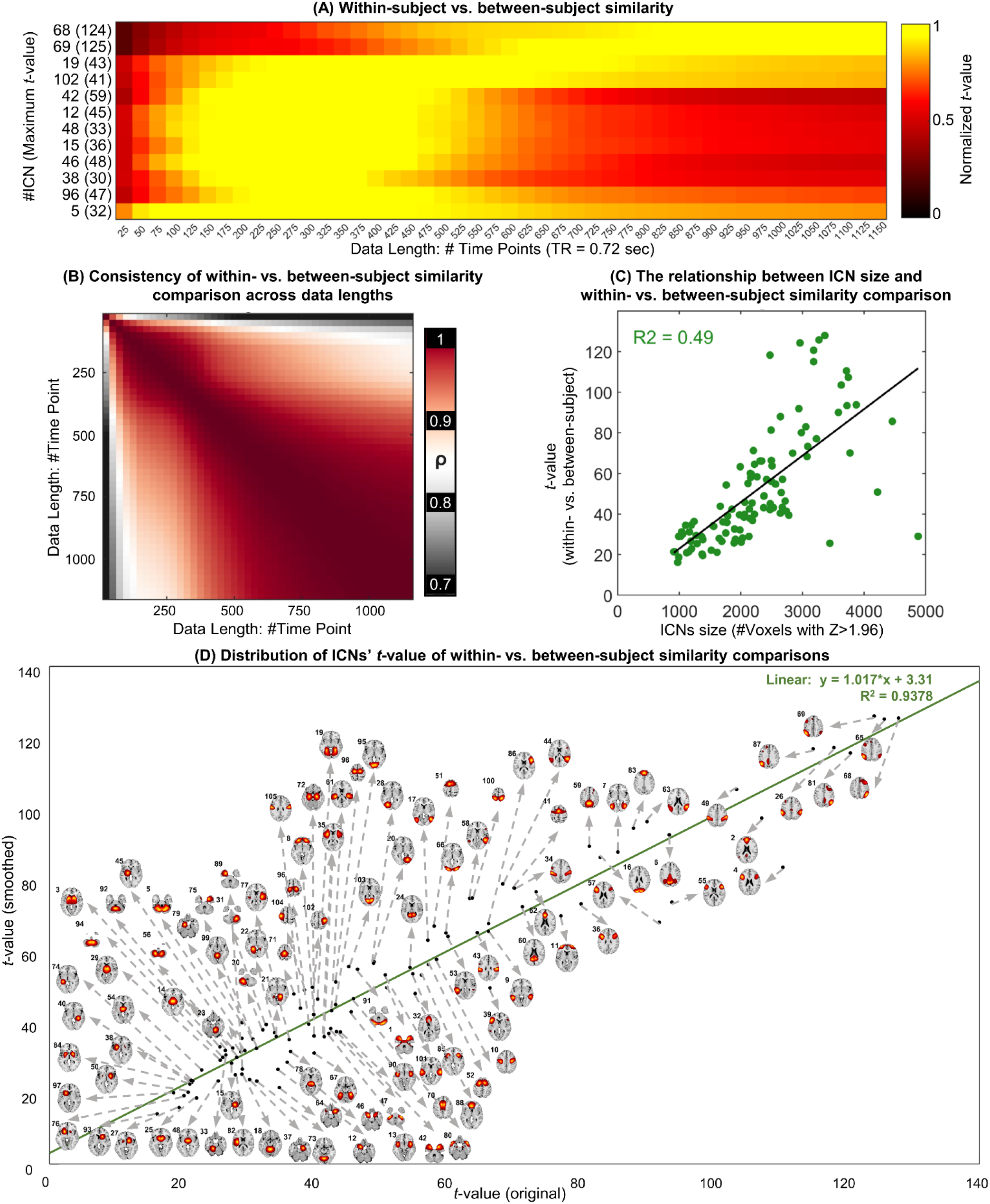
Within-subject versus between-subject comparison. (A) shows examples of this comparison for selected ICNs and different data lengths. (B) illustrates similarity in ICNs’ t-values across different data lengths. (C) the relationship between the t-value of within-versus between-subject comparison and the size of the ICNs. (D) demonstrates that the subject-specificity pattern is similar for original estimations and smoothed ones and the distribution of ICNs’ subject-specificity power.

## 4. Discussion

rsfMRI is a non-invasive brain imaging method with arguably the best existing spatial and temporal resolution trade-off and minimal demand from individuals during data acquisition. These properties make rsfMRI a promising tool for studying brain function and for use in clinical applications. Among different features acquired from rsfMRI, functional connectivity, which assesses the interactions across the brain, has shown associations with various mental and cognitive measures, as well as characteristic alterations in certain brain disorders. However, the limitations of existing methods and datasets prevent us from fully leveraging the potential of rsfMRI to study FC and transition it into a well-established, valid clinical tool.

A key step toward establishing rsfMRI as a prevalent clinical tool is the accurate estimation of corresponding functional patterns, i.e., the identification of equivalent functional patterns across individuals and brain states in a way that captures both individual variations and inter-subject correspondence. Accumulating evidence of spatial differences in functional patterns across individuals and even within individuals over time (Bhinge et al., 2019; Boukhdhir et al., 2021; Fan et al., 2021; Iraji et al., 2019a; Iraji et al., 2019c; Iraji et al., 2020; Luo et al., 2021; Salehi et al., 2020; Wu et al., 2021) highlights the necessity of using data-driven approaches instead of predefined (anatomical or functional) atlases in FC studies. However, several factors must be taken into account when using data-driven approaches. First, the calculation of functional patterns for each subject should preferably be estimated independently from other subjects, particularly in the case of prediction and machine learning, which require the training and testing data to be completely separate. Data-driven approaches that separately estimate functional patterns for each subject and then use a data-driven matching technique to find the correspondence fail to adequately satisfy this criterion because the matching step requires comparing the functional patterns across all data (training and testing) and therefore results in data leakage. Second, differences in datasets across studies can influence results. For example, changes in datasets used in a study (e.g., adding new subject data) can impact the matching steps (and dataset-specific group-level estimates) and lead to different results. Furthermore, we often do not have access to all data used in previous studies, and even if we do, rerunning analyses by adding new data to the previous ones each time is impractical and can lead to different solutions. To address the abovementioned complications, a straightforward and practical solution is to use reliable templates and group-informed data-driven techniques to obtain corresponding functional patterns for each individual separately.

Here, we contribute to the solution above by identifying reliable group-level multi-spatial-scale ICNs (as an estimation of a universal template) using data from over 100k subjects and gr-msICA. We used ICA to obtain our templates because of several key factors. First, ICA is a widely used multivariate tool that divides the brain into temporally coherent patterns, known as intrinsic connectivity networks (ICNs), which are potentially spatially overlapping, yet functionally distinct patterns (Calhoun and Adali, 2012; Calhoun and de Lacy, 2017; Iraji et al., 2022), and therefore good estimates of intrinsic functional “sources” or entities (Iraji et al., 2022; Iraji et al., 2020). Another appealing attribute of ICA is its ability to separate artifactual signals from ICNs in the mixed rsfMRI time series (Calhoun and de Lacy, 2017). As such, FNC estimations (both intra- and inter-network FC) have been shown to be more robust to artifacts and less contaminated with erroneous signals compared to other FC measurements (Calhoun and de Lacy, 2017). Furthermore, an ICN’s spatial map has a value at every voxel, indicating the contribution of each voxel to the ICN. Therefore, instead of splitting the brain into separate parcels, ICA appreciates the brain’s functional heterogeneity and multifunctionality (Calhoun et al., 2009; Haak and Beckmann, 2020; Iraji et al., 2020). Another major advantage of ICA is its ability to capture ICNs across multiple spatial scales without imposing a hard constraint on the spatial extent of ICNs (Iraji et al., 2022). This is an important attribute because the spatial scale of functional systems in the brain does not necessarily change at the same rate. We used gr-msICA (Iraji et al., 2022; Meng et al., 2021) to estimate ICNs across multiple spatial scales and to obtain a more complete view of brain function. Supplementary 4 and 5 contain corresponding ICNs can be obtained from single low and high model orders (25 and 175) for those interested in using single scale analysis at only large spatial scale or fine-grained ones.

Moreover, our results show different ICNs may have different optimal model orders across different datasets. Our results show that, for 28 of the 105 ICNs, the best-matched components came from different model orders across 100 half-split subsets of the QC-passed dataset, and the same was observed for 53 out of 105 ICNs in the QC-failed dataset. These findings demonstrate that gr-msICA can significantly improve the stability of ICA results and the identification of corresponding ICNs across datasets. The ability to identify the same ICNs in new data is challenging for group-level ICA analysis and can impact its replicability, particularly when the datasets are independent and have different characteristics (Du et al., 2020). It should be noted that while our findings show ICNs are identified across different ranges of ICA model orders (see examples of different ranges in Figure 2(D)), we posit all would appear across multiple model orders, with one model order exhibiting the highest stability value (similar to ICN 8 and 68) if we had used a larger range of model orders with smaller intervals (e.g., model order 2 to 500 with an incremental step of 1). It is also worth mentioning that the ICNs can be used to estimate a comprehensive subject-specific (and group-level (Joliot et al., 2015)) canonical parcellation/atlas.

A prior study uses 1005 and 823 typical control individuals from the HCP and the Genomics Superstruct Project (GSP) datasets and group level spatial ICA to obtain a template (Du et al., 2020). Our work is different and improves this initial template in several key aspects. In this study, we used a much larger dataset (over 57k individuals after quality control) from various demographics (not just typical control) to extract our ICN template; as such, it provides a closer estimation of a global template. Another major difference is that, rather than using a single model order of 100, here we leverage gr-msICA to improve the consistency of estimated ICNs and to obtain a more comprehensive template of ICNs across multiple spatial scales. Moreover, we use a much higher spatial correlation value as the threshold (0.8 versus 0.4) for the reproducibility and stability of ICNs. If we were to use a threshold of 0.8, the data from Du et al. template shows only 16 replicable ICNs. In contrast, all 105 ICNs of our template have spatial similarity above 0.8 with an unseen independent QC-failed dataset. We also evaluated the presence of the Du et al. ICNs in our selected ICs. For the threshold of 0.4, we found correspondence for all 53 template ICNs among 900 ICs. However, only 45 out of 53 ICNs for Du et al., show spatial similarity above 0.4 with our 105 template ICNs; in other words, some of the ICNs in the initial template did not meet our criteria of being an ICN.

Focusing on sample-specific (e.g., subject-specific) estimation, we used group-informed network estimation techniques where our template was used to guide subject-specific solutions. We recommend using group-informed network estimation techniques combined with our template for several reasons. Firstly, group-informed techniques more accurately estimate subject-specific patterns, particularly for low SNR rsfMRI data with short data lengths, because the templates are utilized as a constraint to limit the search space. A recent study (Duda et al., 2022) suggests that this pipeline could potentially shorten clinical rsfMRI scans to just 2-4 minutes without significant loss in static group comparison. Group-informed techniques also enhance the estimation of subject-specific correspondence by optimizing the solution to be jointly spatially independent and close to a common template. In addition, these techniques only use the template and subject data itself to estimate subject-specific ICNs, and therefore the estimation of each individual is independent of other samples in a given study. Using our template as a universal reference also facilitates the comparison of findings across existing and future studies as we retain the correspondence ICNs across all subjects that the pipeline is applied to.

The first step toward using any subject-level estimates is to evaluate whether or not network-estimation and parcellation techniques successfully estimate subject-level patterns for a given dataset. This has been mainly overlooked in previous studies. Here, we introduced two criteria for this purpose which assess whether the subject-level estimates of each ICN in a given dataset provide (1) subject-specific information beyond predefined spatially fixed nodes (weighted masks) and (2) unique information about the associated ICN compared to other ICNs. The second criterion is important for highly similar ICNs and evaluating if we can differentiate between them in subject-level estimation in a given dataset. In this work, we assessed the ability of the existing MOO-ICAR framework to obtain the subject-level estimation of our template. But these two criteria can be used to evaluate other network-estimation and parcellation techniques, and we also call for further investigation on this understudied but important area.

We also evaluated the role of different parameters in estimating subject-level ICNs. The results show that in addition to the data length, which has been the center of many reliability investigations (Birn et al., 2013; Duda et al., 2022; Gordon et al., 2017b; Murphy et al., 2007), other data characteristics (particularly inherent spatial resolution) play key roles in successfully extracting subject-level ICNs. For instance, MOO-ICAR successfully estimated all 105 ICNs using 100 time points for the HCP dataset; however, 12 ICNs did not survive the second criterion for the FBIRN dataset, even using 157 time points. This might suggest the spatial resolution of data is an important factor in differentiating these highly spatially similar ICNs. These highly spatially similar ICNs (Supplementary 6) that we, for the first time, observed at the group-level analysis resemble the previously reported parallel interdigitated distributed networks observed at the subject level (Braga and Buckner, 2017).

In addition to data characteristics, the intrinsic properties of a given ICN are an important factor in calculating subject-level estimates. In general, larger ICNs require less data to achieve a specific level of within-subject (between-subject and template) spatial similarity. Given that spatial similarity is commonly used as a reliability index, we could postulate large-scale ICNs are more stable and requires less data to be estimated reliably at the subject level. This result agrees with previous findings showing that lower model-order ICA generates more consistent components than the higher model orders. Our findings also indicate that ICNs carry different levels of subject fingerprint information, with ICNs associated with the subcortical domain having the least subject-specific information and those suggested to be involved in higher cognitive functions, particularly ICNs associated with frontoparietal domains showing the maximum within-to-between-subject differences. A previous fingerprint study also identified the connectivity patterns of the frontoparietal network as the most distinguishing of individuals (Finn et al., 2015).

Interestingly, ICNs exhibit different patterns of within-to-between-subject differences across data lengths, which can relate to the temporality aspect of functional fingerprinting (Van De Ville et al., 2021), for example, the within-to-between-subject difference peaks at different data lengths for different ICNs. Our analysis using the HCP dataset with TR = 0.72 seconds and a step size of 25 time points shows the peak varies between 125 to 875 time points (90 to 630 seconds). These findings may also imply (1) increasing data length is not always desirable even though it increases within-subject similarity, which is commonly used in reliability analyses to support using longer data lengths (30 mins and more) in analysis, (2) studies should take the length of the data into account in their analyses and interpreting the results, and (3) future work can analyze and leverage multi-data-length information. One related key factor that was not explored in this study is incorporating and assessing the brain dynamics. The lower spatial similarity for smaller data lengths can be partially related to brain spatial dynamics and changes in the spatial patterns of ICNs over time (Iraji et al., 2019a). These differences observed in findings across ICNs and data lengths (and probably other factors) highlight the challenge of within- and between-subject variabilities in understanding brain functional organization and its changes across different conditions.

Notably, while there are observable differences in findings between data lengths, the findings across various data lengths still show a similar pattern across ICNs. For instance, while within-subject and between-subject similarities are different across data lengths (i.e., a systematic increase in spatial similarity as a function of data length), the pattern of spatial similarity across ICNs remain fairly similar across data lengths (e.g., Figure 5(C) and (D)), suggesting features that encode ICN properties relative to each other might be more robust to the data length and therefore possibly more generalizable indicators of brain function.

We also observe that differences in spatial smoothness across data lengths (also between datasets) can impact the results and outcome of analyses, and therefore may limit the comparison of findings across studies. We show how spatial smoothing can alter results, including improving within-subject spatial similarity. As such, we highlight the necessity of correcting for differences in spatial smoothness, particularly those associated with data lengths, for any interpretation of results and comparing the findings across analyses.

Finally, the within-session results (i.e., our analyses of different overlapping data within each session for different data lengths) may indicate that the lower spatial similarity in smaller data lengths may not be solely related to lower reliability in estimation but also associated with the dynamic nature of brain function. We observed that the within-session similarity between ICNs estimated using the full-length of data and subset of it (e.g., 50% and 25% data lengths) remains fairly similar for different full-lengths of data (cf. blue and green curves are relatively flat in Figure 5(H)). For example, the spatial similarity between ICNs estimated using 100 time points and the first 50 time points is close to 0.8, as is the similarity between 1000 time points and the first 500 time points. We would expect a significantly lower spatial similarity between 100 and 50 time points if their ICNs estimates were unreliable compared to those from longer data lengths. We posit that the lower between-session spatial similarity for lower data lengths might be because the two separated data carry different spatial dynamic information of ICNs, and as the data lengths increase, they are gradually getting closer to average (also known as static) estimates of ICNs, and therefore become more similar. Indeed, the larger increase in the spatial similarity of between-session analysis relative to within-session analysis may further support this posit.

## 5. Conclusion

In this work, we identified a reliable and replicable multi-spatial-scale ICNs template using gr-msICA and around 58k subject data that meet the quality control criteria. The template was also replicated in data that did not pass the QC criteria. We aim to use this template to generalize and standardize functional connectivity analysis. This study builds on the recently proposed concept of Neuromark. NeuroMark is a comprehensive mapping of (unimodal or multimodal) coherent brain patterns which correspond among individuals by leveraging universal templates derived from prior data coupled with guided data-driven approaches. Previously, NeuroMark_fMRI_1.0 template, including 53 ICNs, was obtained from two large rsfMRI datasets and single model order ICA (Du et al., 2020). Here, we augmented the previous effort by using much larger datasets and gr-msICA.

In addition to providing an enhanced ICN template, we also studied the feasibility of estimating the corresponding ICNs at subject-level. Previous work showed additional flexibility and robustness Neuromark framework even across different processing pipelines (DeRamus et al., 2021). But there is a significant gap in evaluating factors that influence successful captures of subject-specific information (ICNs or functional parcellations), due in part to a lack of known ground truth for evaluation of estimates. Here, we introduced two criteria to evaluate the successful identification of subject-specific ICNs (or other functional parcellations) for a given dataset and a group-informed estimation approach and studied the role of different factors in subject-level ICN estimates. The results suggest that intrinsic properties of the ICNs themselves, data length, and spatial resolution are some key factors in successfully estimating ICNs at the subject level. We illustrated an increase in spatial smoothness as a function of data length and the impact of spatial smoothing on findings. As such, we suggest future studies should control for the effect of spatial smoothness in their analysis to mitigate its impact on our ability to compare the findings across different studies. We also observed increasing data length can reduce an ICN’s subject-level specificity, suggesting longer scans might not always be desirable. Finally, the consistency in the spatial similarity between ICNs estimated using the full-length of data and subset of it across different data lengths may suggest that the lower within-subject spatial similarity in shorter data lengths is not necessarily defined by only lower reliability in ICNs estimates and demands further investigations. Our future work will focus on incorporating the findings of this study in functional connectivity analysis and developing new group-informed network estimation techniques to improve the estimation of corresponding subject-specific ICNs. Future research can benefit from using higher model order ICAs and lower step sizes. Future work can explore using other multi-model-order ICA approaches (Du et al., 2021) and develop more advanced gr-msICA to estimate ICNs across multiple spatial scales. We will also soon release the Neuromark templates for other imaging modalities.

## 6. Acknowledgment

This work was supported by grants from the National Institutes of Health grant numbers 1U24RR021992, 1U24RR025736, R01EB020407, R01MH118695, R01MH123610, and R01EB006841 and the National Science Foundation grant number 2112455 to Dr. Vince D. Calhoun; the National Institutes of Health grant number R01MH117107 to Dr. Jing Sui; the National Science Foundation grant number 1631838 to Dr. Tulay Adali; and the Lawson Health Research Institute, grant number LHR D1374, Pfizer Independent Investigator Award, grant number WS2249136, and CIHR grant FRN 153359 to Dr. Elizabeth A. Osuch. We would also like to acknowledge Georgia State University RISE Award, which significantly helped to accomplish this work. Data were provided in part by the Adolescent Brain Cognitive Development SM (ABCD) Study (Jernigan and Brown, 2018), held in the NIMH Data Archive (NDA). This is a multisite, longitudinal study designed to recruit more than 10,000 children age 9–10 and follow them over 10 years into early adulthood. The ABCD Study^®^ is supported by the National Institutes of Health (NIH) and additional federal partners under award numbers U01DA041048, U01DA050989, U01DA051016, U01DA041022, U01DA051018, U01DA051037, U01DA050987, U01DA041174, U01DA041106, U01DA041117, U01DA041028, U01DA041134, U01DA050988, U01DA051039, U01DA041156, U01DA041025, U01DA041120, U01DA051038, U01DA041148, U01DA041093, U01DA041089, U24DA041123, U24DA041147; by the Autism Brain Imaging Data Exchange (ABIDE) (Di Martino et al., 2017; Di Martino et al., 2014) support for the work by Adriana Di Martino provided by the (NIMH K23MH087770) and the Leon Levy Foundation and primary support for the work by Michael P. Milham and the INDI team was provided by gifts from Joseph P. Healy and the Stavros Niarchos Foundation to the Child Mind Institute, as well as by an NIMH award to MPM (NIMH R03MH096321); by the Attention Deficit Hyperactivity Disorder-200 (ADHD200) Consortium (HD-200 Consortium, 2012). Consortium steering committee includes Jan Buitelaar, M.D., F. Xavier Castellanos, M.D., Ph.D., Daniel Dickstein, Ph.D., Damien Fair, P.A.-C, Ph.D., David Kennedy, Ph.D., Beatric Luna, Ph.D., Michael P. Milham (Project Coordinator), M.D., Ph.D., Stewart Mostofsky, M.D., Joel Nigg, Ph.D, Julie B. Schweitzer, Ph.D., Katerina Velanova, Ph.D., Yu-Feng Wang, M.D., Ph.D., Yu-Feng Zang, M.D.; by the Alzheimer’s Disease Neuroimaging Initiative (ADNI) (Jack et al., 2008) (National Institutes of Health Grant U01 AG024904) and DOD ADNI (Department of Defense award number W81XWH-12-2-0012). ADNI is funded by the National Institute on Aging, the National Institute of Biomedical Imaging and Bioengineering, and through generous contributions from the following: AbbVie, Alzheimer’s Association; Alzheimer’s Drug Discovery Foundation; Araclon Biotech; BioClinica, Inc.; Biogen; Bristol-Myers Squibb Company; CereSpir, Inc.; Cogstate; Eisai Inc.; Elan Pharmaceuticals, Inc.; Eli Lilly and Company; EuroImmun; F. Hoffmann-La Roche Ltd and its affiliated company Genentech, Inc.; Fujirebio; GE Healthcare; IXICO Ltd.; Janssen Alzheimer Immunotherapy Research & Development, LLC.; Johnson & Johnson Pharmaceutical Research & Development LLC.; Lumosity; Lundbeck; Merck & Co., Inc.; Meso Scale Diagnostics, LLC.; NeuroRx Research; Neurotrack Technologies; Novartis Pharmaceuticals Corporation; Pfizer Inc.; Piramal Imaging; Servier; Takeda Pharmaceutical Company; and Transition Therapeutics. The Canadian Institutes of Health Research is providing funds to support ADNI clinical sites in Canada. Private sector contributions are facilitated by the Foundation for the National Institutes of Health. The grantee organization is the Northern California Institute for Research and Education, and the study is coordinated by the Alzheimer’s Therapeutic Research Institute at the University of Southern California. ADNI data are disseminated by the Laboratory for Neuro Imaging at the University of Southern California; by the Bipolar & Schizophrenia Consortium for Parsing Intermediate Phenotypes (B-SNIP) study (Tamminga et al., 2013); by the Brain Genomics Superstruct Project (**GSP**) (Holmes et al., 2015) of Harvard University and the Massachusetts General Hospital, (Principal Investigators: Randy Buckner, Joshua Roffman, and Jordan Smoller), with support from the Center for Brain Science Neuroinformatics Research Group, the Athinoula A. Martinos Center for Biomedical Imaging, and the Center for Human Genetic Research. 20 individual investigators at Harvard and MGH generously contributed data to the overall project; by the Human Connectome Project for Early Psychosis study (Lewandowski et al., 2020); the Human Connectome Project (Van Essen et al., 2013), WU-Minn Consortium (Principal Investigators: David Van Essen and Kamil Ugurbil; 1U54MH091657) funded by the 16 NIH Institutes and Centers that support the NIH Blueprint for Neuroscience Research; and by the McDonnell Center for Systems Neuroscience at Washington University; by the OASIS (LaMontagne et al., 2019) Longitudinal Multimodal Neuroimaging: Principal Investigators: T. Benzinger, D. Marcus, J. Morris; NIH P50 AG00561, P30 NS09857781, P01 AG026276, P01 AG003991, R01 AG043434, UL1 TR000448, R01 EB009352. AV-45 doses were provided by Avid Radiopharmaceuticals, a wholly owned subsidiary of Eli Lilly; and using the UK Biobank (Littlejohns et al., 2020) Resource under Application Number 49636.

